# A multi-flow approach for binning circular plasmids from short-reads assembly graphs

**DOI:** 10.64898/2026.03.25.714305

**Authors:** Victor Epain, Aniket Mane, Gianluca Della Vedova, Paola Bonizzoni, Cedric Chauve

## Abstract

We address the problem of plasmid binning, that aims to group contigs – from a draft short-read assembly for a bacterial sample – into bins each expected to correspond to a plasmid present in the sequenced bacterial genome. We formulate the plasmid binning problem as a network multi-flow problem in the assembly graph and describe a Mixed-Integer Linear Program to solve it. We compare our new method, `PlasBin-HMF`, with state-of-the-art methods,`MOB-recon, gplasCC`, and `PlasBin-flow`, on a dataset of more than 500 bacterial samples, and show that `PlasBin-HMF` outperforms the other methods, by preserving the explainability.

## 1 Introduction

Plasmids are extra-chromosomal Mobile Genetic Elements (MGE) present in bacterial genomes that can easily transfer between bacterial cells, possibly of different bacterial species, and contribute widely to the propagation of genes responsible for Antimicrobial Resistance (AMR) [15,6]. Due to the importance of AMR in public health [18], an important aspect of epidemiological surveillance focuses on detecting plasmids in bacterial sequencing data [5]. Short-read sequencing is still the most commonly used data acquisition technology for epidemiological surveillance. As a consequence, the development of bioinformatics methods for the detection and characterization of plasmids in bacterial draft assemblies obtained from short-read sequencing data is an active research area [1,17,3,16,13,19,23,10,20,14].

Detecting plasmids from a short-read draft assembly can be addressed at three different levels: contigs classification, plasmid binning and plasmid assembly. *Contigs classification* aims to separate contigs between plasmidic contigs and chromosomal contigs, although some contigs can also be classified as ambiguous if they are repeats present both in the chromosome and in some plasmid. There is a large corpus of classification methods, most of them based on machine-learning approaches [22]. As a bacterial genome can contain several plasmids, contigs classification does not provide a precise view of its plasmid content. This is addressed by *plasmid binning*, that computes groups of contigs (*plasmid bins*) with the aim that each plasmid bin contains the contigs constituting a single plasmid. *Plasmid assembly* can be seen as a refinement of plasmid binning where contigs in each plasmid bin are ordered and oriented.

`MOB-recon` [16] is one of the most popular plasmid binning methods, based on aligning contigs to a database of known plasmid-specific genes and a curated database of complete plasmid sequences. To the best of our knowledge, all other plasmid binning methods rely on a different approach that makes use the of *assembly graph*, a graph provided together with contigs by assemblers widely used for bacterial data, `SPAdes` [4], `Unicycler` [24] and `SKESA` [21]. The first methods of this kind were `plasmidSPAdes` [1] and `Recycler` [17]. Both rely on the expectation that contigs from the same plasmid (forming a true plasmid bin) should be co-located in the assembly graph and should have read coverage that is different from the expected coverage of chromosomal contigs and uniform within a plasmid bin. Unlike `MOB-recon`, `plasmidSPAdes` and `Recycler` are *de novo* methods that do not rely on homology with known plasmids. `HyAsP` [13] introduced the notion of scoring plasmids bins based on several features including the coverage of contigs by known plasmid genes, the uniformity of read coverage and of the GC content, and integrated this scoring scheme in an iterative greedy heuristic searching for high-score walks in the assembly graph, each such path forming a plasmid bin. `gplas` [2], similarly to `HyAsP`, is a greedy walk detection heuristic but it relies on a different scoring function and introduced the idea of using plasmid contigs classification results as an important signal to detect plasmid bins; the latest development of this method is `gplasCC` [14]. The `HyAsP` scoring function served as a basis for `PlasBin` [9], the first approach that introduced a rigorous combinatorial optimization approach to plasmid binning; `PlasBin` iteratively detected the plasmid bin of maximum score using a Mixed-Integer Linear Program (MILP), peeled the corresponding subgraph out of the assembly graph, and repeated until no good plasmid bin could be found. `PlasBin-flow` [10] improved upon `PlasBin` by using the concept of network flow to define plasmid bins as connected subgraphs that would optimize a function combining the features used in `PlasBin` and the flow value, considered as a proxy for the copy number of the hypothetically detected plasmid; however, similarly all other graph-based methods discussed above, `PlasBin-flow` is an iterative method, searching for one plasmid bin at a time.

Here we propose a novel plasmid binning method, `PlasBin-HMF` (standing for Plasmid Binning Hierarchical Multi-Flow), that builds upon the high-level principles of `PlasBin-flow` but does not search for plasmid bins one at a time and, instead, relies on the concept of multi-flow to detect a set of plasmid bins at once. To the best of our knowledge, `PlasBin-HMF` is the first plasmid binning method that defines the problem rigorously from a combinatorial optimization point of view and solves it exactly using an MILP. In Section 2, we formulate the plasmid binning problem as a multi-flow combinatorial optimization problem and describes how it can be solved using an MILP. In Section 3 we apply `MOB-recon`, `gplasCC`, `PlasBin-flow` and `PlasBin-HMF` to a dataset of more than 500 bacterial samples, and we show that `PlasBin-HMF` outperforms the other methods. We conclude by discussing avenues for improving the multi-flow model that serves as a basis for `PlasBin-HMF` .

## 2 Methods

Informally, given sequencing data from a bacterial sample and its assembly, plasmid binning aims to determine the contig content (a set of contigs) of all plasmids in the sample, without prior knowledge about the number of plasmids or their genomic sequences. From now on, we refer to a set of contigs as a *bin*. In this section we describe a combinatorial optimization model to detect plasmid bins; we focus on describing the mathematical model, and postpone the description of the MILP formulation to the Supplementay Material. In the description that follows, we assume that plasmids are circular molecules, which is the case in the vast majority of cases.

### 2.1 Input data

#### Assembly graph

The assembly of short-read sequencing data results in a (unoriented) contig set 𝒞 and a link set ℒ, a link being an ordered pairs of oriented contigs, ℒ ⊂ (𝒞 *× {*+, −*}*)^2^, where + and − respectively stand for the forward and the reverse orientations. Note that for each link (*c, d*) ∈ℒ, its reverse 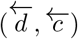 is also in ℒ, where 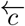 and 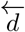 the reverse of the oriented contigs *c* and *d*. The link set naturally defines a directed graph, called the *assembly graph*, where every vertex corresponds to an oriented contig (see Figure S1 in Appendix A).

#### Contigs length and coverage

For a contig *c* ∈𝒞, we denote by len_*c*_ its length and by cov_*c*_ its *normalized coverage*, i.e. its average read coverage divided by the mean read coverage over all contigs. The normalized coverage serves as a proxy for the expected number of copies of the contig sequence in the sequenced genome. The widely used assembler Unicycler provides this information by default.

#### Plasmidness score and plasmid seeds

We also assume that each contig is provided with a *plasmidness score* plm_*c*_ [− 1, 1] that estimates the nature of the contig as either chromosomal (plm_*c*_ *<* 0) or plasmid-derived (plasmidic) (plm_*c*_ *>* 0). Accurate plasmidness scores would assign a score of 1 to purely *chromosomal* contigs, 1 to purely *plasmidic* contigs, and 0 to contigs appearing both in the chromosome and in some plasmid(s) (*ambiguous* contigs). Finally, we assume that we are provided with a subset 𝒞 ⊂ 𝒞 _*seed*_ of *seed* contigs, which are estimated to be plasmid-derived with high confidence. We describe in Section 3 how we use the plasmid contigs classifications tools `RFPlasmid` [23], `plasmidCC` [14] and `Platon` [19] to compute plasmidness scores and determine plasmid seeds.

### 2.2 Overview

Most plasmid binning methods that rely on exploring the assembly graph are based on the following principle: if we assume (1) uniform sequencing depth, (2) an accurate assembly graph, and (3) an accurate classification of the contigs as chromosomal, plasmidic or ambiguous, then a circular plasmid appears in the assembly graph as a circuit composed of plasmidic (and possibly ambiguous) contigs, with uniform normalized coverage that approximates well the copy number of the plasmid. The contig content of this circuit forms the (true) plasmid bin for the corresponding plasmid.

`PlasBin-HMF` is based on the same high-level principle, accounting for expected inaccuracies in the input data. Indeed, in practice, (1) sequencing depth is not uniform, (2) assembly graphs may exhibit spurious or missing links, and (3) contigs classification methods are prone to errors. `PlasBin-HMF` allows to define a (plasmid) bin as a connected subgraph of the assembly graph (instead of a circuit, to handle point (2)), whose contigs’ normalized coverage is consistent with a flow through this subgraph (point (1)). `PlasBin-HMF` allows a bin to contain contigs classified as chromosomal but integrates in its scoring function the plasmidness score such that it assigns a better score to subgraphs that are denser in plasmidic contigs (point (3)). Unlike iterative methods, one of the main novelties of `PlasBin-HMF` is that it simultaneously identifies in the assembly graph multiple subgraphs and flows, each corresponding to a single plasmid bin.

### 2.3 Multi-flow modelling of plasmid bins

We first describe the network derived from the assembly graph, on which we model plasmid bins as network flows (Definition 1). As we potentially search for several bins, we model one flow per bin (Definition 2, Figure 1). We focus our search to circular bins in the network (Definition 3), and complete the list of bins with partially circular ones in the case some links are missing (Definition 4). We then model the plasmid binning as a combinatorial optimization problem on the network, using a multi-flow formulation framework, that we formalize as an MILP (Appendix B.2).

**Fig. 1.**
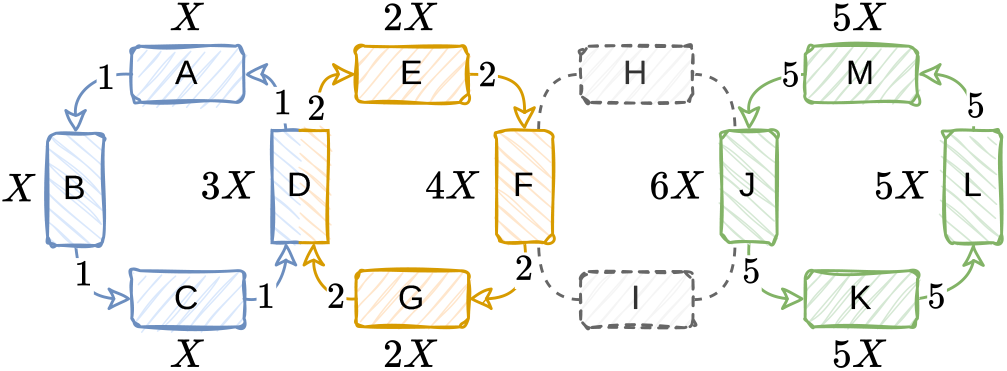
Example of an assembly graph with three circular plasmid bins, {A, B, C, D },{D, E, F, G} and {J, K, L, M}, with respective copy numbers 1, 2 and 5. The contig coverages are written as *kX, k* ∈ℕ. The positive flows are numbers on the arcs. Contig D is shared between the blue and the orange plasmids. The total incoming flow in D equals its coverage (1 + 2 = 3). The coverages of F and J are not fully explained by the flows, so one could assume they also belong to other elements of the genome than plasmids.

#### Definition 1

**(Flow network)**. *Let G* = (*V, A*, s, t) *be the flow network defined as follows:*

– *the set V of vertices contains the vertices of the assembly graph (associated to the oriented contigs), the source* s *and the sink* t: *V* = *V*_𝒞_ ∪ {s, t}, *where V*𝒞 =𝒞 *×* {+,*−* };^1^
– *the set A of arcs contains the link-arcs of the assembly graph and the arcs from the source and to the sink: A* = ℒ ∪ *{*(s, *v*) | *v* ∈ *V*_𝒞_*}* ∪ *{*(*v*, t) | *v* ∈ *V*_𝒞_*};*^1^

Retrieving the contig from a vertex and conversely is enabled thanks to two functions *vtoc* : *V*_*𝒞*_ →𝒞 and 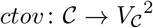. For each vertex *v, V*^−^(*v*) and *V* ^+^(*v*) denote respectively the set of predecessors and successors of *v* in the network. For a vertex *v* ∈ *V*, 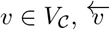 denotes its reverse (the same contig but with a reversed orientation, so 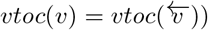

#### Definition 2

**(Coverage-constrained multi-flow)**. *A multi-flow set of size n* ∈ ℕ_≥1_ *is a set of n functions {f*_1_, …, *f*_*n*_*} such that:*

**–** ∀1 ≤ *k* ≤ *n, f*_*k*_ : *A* → R_≥0_

**–** ∀*c* ∈ 𝒞:

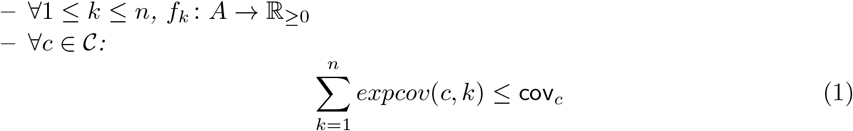

*where*

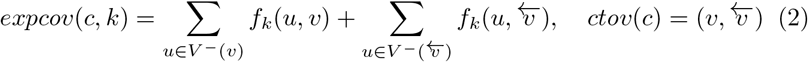

**–** ∀*v* ∈ *V*_*C*_, 1 ≤ *k* ≤ *n:*

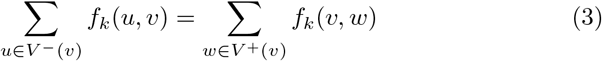

**–** ∀1 ≤ *k* ≤ *n:*

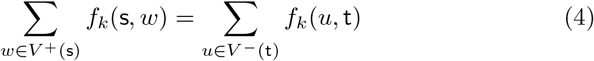

#### Modelling plasmids as multi-flows

As discussed in Section 2.2, a bin is the set of contigs of a subgraph of the assembly graph. In `PlasBin-HMF`, each flow in a multi-flow set induces a non-empty subgraph, and thus a bin. We first search for a multi-flow composed only of circular flows (Definition 3), that models putative plasmids that appear as circuits in the assembly graph. Then we relax the circularity constraint (Definition 4) to account for possibly missing links in the assembly graph. Figure 2 illustrates the two kinds of structures we are looking for.

**Fig. 2.**
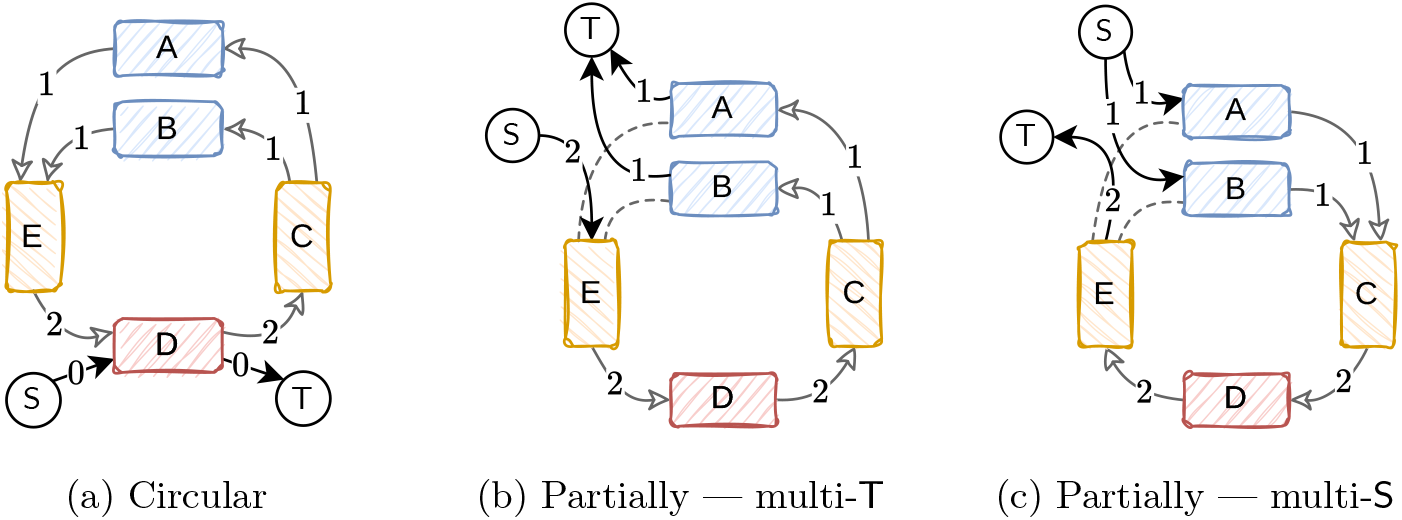
Fully and partially circular flows. In each subfigure, the red contig D is a seed, orange and red contigs have a coverage of 2*X* while the blue contigs A and B have coverage *X*. The numbers on the arcs are the flow values. For the ease of read, we do not show other links between other plasmids or chromosomes and suppose we cannot merge C, D and E in one unitig. The links are oriented according to the orientation of the positive flow. **(a)** No link is missing such that we can find a circuit using all the contig coverages. **(b)** — **(c)** Dashed links are missing links. The source S and the sink T simulate the circularity by connecting the extremities of the partial circuit.

##### Definition 3

**(Circular coverage-constrained flow)**. *A function f in the coverage constrained multi-flow set is circular if the set of strictly positive flow links {a* ∈ ℒ | *f* (*a*) *>* 0*} is defining a circuit in G*.

##### Definition 4

**Definition 4 (Partially circular coverage-constrained flow)**. *A function f in the coverage constrained multi-flow set is partially circular if:*

– *the set of strictly positive flow arcs augmented with an artificial arc from the sink to the source* {*a* ∈ *A* | *f* (*a*) *>* 0} ∪ (t, s) *is defining a circuit in G;*
– *the subgraph without* s *and* t *is connected;*
– *each vertex connecting the source, respectively the sink, has no other positive-flow predecessor, resp. successor, i*.*e*.

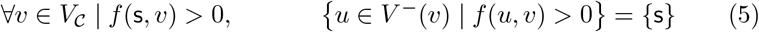

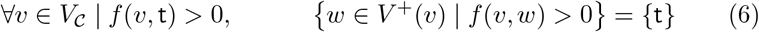

Definition 4 ensures the contigs connected by the source and the sink are extremities of the “cut circuit”. It prevents to artificially connecting contigs from other plasmids or misidentified non-plasmidic contigs to the extremities of the cut circuit.

Last, we describe how we make use of the plasmidness score of contigs in defining plasmid bins. Each flow function *f*_*k*_ in the multi-flow set is associated to a plasmidness score (Definition 5). The more the flow uses the coverage of a long plasmidic (non-plasmidic) contig, the better (worse) the score.

##### Definition 5

**Definition 5 (Plasmidness score)**. *The plasmidness score of a flow function f*_*k*_ *in a multi-flow set is defined as:*

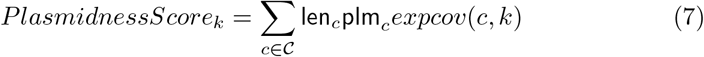

In what follows, we require that any flow that would define a plasmid bin has a positive plasmidness score.

#### Scoring a multi-flow set

We describe how we associate with a multi-flow set a linear *explanation score* (Definition 6), where a higher scores implies a better explanation of the bins defined by the flows in terms of the input data. `PlasBin-HMF` aims to search for a multi-flow set maximizing the explanation score. On one hand, we favour multi-flow sets that maximizes the use of plasmidic contigs, avoiding the ones which are not (*PlasmidnessScores* term). On the other hand, we penalize multi-flow sets when they fail to fully explain the coverages of plasmidic contigs (*PosPlmCovPenalty* term). We model the search of the best multi-flow set of size *n* through an MILP that we describe formally in Appendix B.2.

##### Definition 6

**(Explanation score)**. *Given a multi-flow set of size n, its explanation score is defined as follows:*

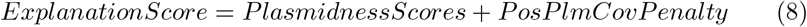

*where:*

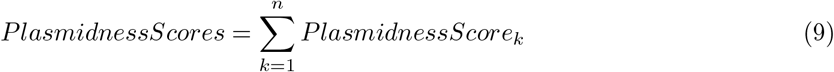

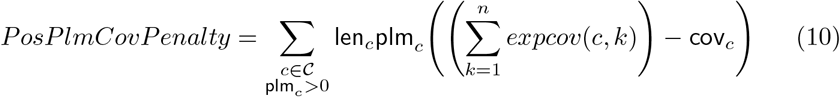

#### Deciding the number of flows

In what precedes, we always assumed that the number of flows in a multi-flow set was given. As the number of plasmid bins is exactly the number of flows in a multi-flow set, the question of choosing the size (number of flows) that is searched by `PlasBin-HMF` is an important question. Informally, we address this problem by computing optimal multi-flow sets of increasing size until a parsimony stopping criterion, based on the comparison among the explanation scores of successive multi-flow sets, is met. Assume that during this process, at iteration *m*, we aim to find the best multi-flow set of size *m*. Each time we increment the number of flows, we penalize the next explanation score with an *additional flow penalty* (Definition 7). Given two parameters <u>L</u> (minimum length of a plasmid bin) and <u>cov</u> (minimum flow allowed to define a plasmid bin), the additional flow penalty is the plasmidness score of a flow defining a plasmid bin with a single contig, of length <u>L</u>, plasmidness of 1 and normalized coverage <u>cov</u>. We stop incrementing the number of flows if (1) either we reach the maximum number of flows *n* ∈ ℕ _≥1_ we allowed to search for (see next paragraph) (2) the search is infeasible (3) or the new explanation score plus the additional flow penalty is not (strictly) greater than the previous explanation score (if *ExplanationScore*_*m*+1_ − FlowPenalty ≤ *ExplanationScore*_*m*_, 1 ≤ *m < n*).

##### Definition 7

**(Additional flow penalty)**. *Given two parameters* <u>L</u> ∈ ℕ *and* <u>cov</u> ∈ ℝ_*>*0_ *the additional flow penalty is defined by:*

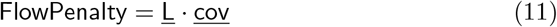

#### Search structure and parameters

A multi-flow set is associated with a plasmid structure (fully or partially circular), which must be described by its flows. Additionally to the subgraph topology, the multi-flow set must respect a seed constraint: we first search for subgraphs containing at least one seed contig, and then relax the constraint. We first search for circular bins with seeds and then without seeds, and repeat the same procedure for partially circular bins, first with seeds and then without. The seed constraint provides us the maximum number of flows a multi-flow set can contain. When each bin must contain a seed, the maximum number of flows is the number of seeds (*n* =| 𝒞_*seed*_ |). Otherwise, the upper bound equals to a user parameter 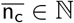, respectively 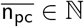 when we search for circular flows, resp. for partially circular flows. Algorithms B.1 and B.2 detail the search prioritisation.

Finally, given a multi-flow set with a circularity and a seed constraints, we require that each flow function *f*_*k*_ (1) has a positive plasmidness score (Definition 5), (2) equals at least the normalized coverage parameter cov when it positively passes through an arc (i.e. *f*_*k*_ : *A* → {0} ⋃ [<u>cov</u>, *∞*]), and (3) describes a bin such that the sum of its contigs lengths at least equals the parameter <u>L</u>.

## 3 Results and Discussion

We implemented the method in `Python3` as a submodule of a larger package https://github.com/AlgoLab/pangebin^2^. We evaluated `PlasBin-HMF` against `MOB-recon`, `PlasBin-flow` and `gplasCC` on 581 bacterial samples for which both short-read and long-read are publicly available on NCBI. In particular, long-read data was used only to define the ground truth plasmid bins. We detail in Appendix C how we obtained these 581 samples.

### 3.1 Data preparation and experiments

#### Input data preparation

For each sample, we assembled the Illumina short reads with `Unicycler`, a read assembler aiming the completion of the circularity. Contigs less than 100bp were removed from the assembly graph and transformed into as many links required to connect their neighbours. The plasmidness of contigs was computed with the classification tools `RFPlasmid` . The plasmidness in `PlasBin-flow` are between 0 and 1, while those of `PlasBin-HMF` are between *−* 1 and 1 (affine transformation). Internally, `PlasBin-HMF` raises the plasmidness of the seed contigs by computing the mean between one and the original input plasmidness. The seeds for `PlasBin-flow` and `PlasBin-HMF` are the contigs identified as plasmid contigs by `Platon` (binary classification).

#### Ground truth

For each sample, we produced a hybrid assembly using both the short and the (Oxford Nanopore) long reads. If a hybrid contig was longer than 1Mbp, we annotated it as chromosomal, if it is circular and less than 1Mbp we considered the contig to be a plasmid, otherwise we declared the contig unlabelled. Here, all the 581 samples have a ground truth with hybrid contigs labelled as plasmids or chromosomal, and no contig is unlabelled. For each sample, we mapped the short contigs against the hybrid ones with `minimap2` [8]. We labeled a short contig as plasmidic if it mapped to a plasmidic hybrid contig, and as non-plasmidic otherwise (i.e., if it mapped only to chromosomal hybrid contigs or to no hybrid contig). To each plasmidic hybrid contig corresponds a ground truth bin, containing the plasmidic short contig that mapped to the hybrid contig. Note that, contrary to the recent plasmid binning method benchmark study [22], a short contig can belong to multiple bins in our case.

#### Experiments

We use the default option values for `MOB-recon`, `PlasBin-flow` and `gplasCC` . For `PlasBin-HMF`, we searched for fully and partially circular bins with seeds, and complete the results with no more than two circular bins without seeds. We also fixed the minimum cumulative contig length a bin reach to <u>L</u> = 1000, and a minimum of flow <u>cov</u> = 0.3. As `gplasCC` outputs bins that exclusively contain contigs classified as plasmidic (even if chromosomal classified contigs participate in the walks connecting the plasmidic ones) we add to the comparison a modified version of `PlasBin-flow` and of `PlasBin-HMF` that mimic the `gplasCC` filtering behaviour. Thus, for each bin, we only keep its seeds and its contigs with a plasmidness greater than 0.5 for `PlasBin-flow` and greater than 0 for `PlasBin-HMF` .

We ran the tools and `PlasEval` on the Digital Alliance Canada high-performance computing clusters Cedar and Fir. Each sample was processed using with 16 CPUs, 32GB of RAM, with a time limit set to 10 hours. `PlasBin-flow` and `PlasBin-HMF` ran with the MILP solver Gurobi.

### 3.2 Accuracy Measures

We evaluate the methods according to two accuracy measures. The first has been introduced in `PlasBin-flow` [10] and is an adaptation of the recall, precision and the F1 statistics, weighted by contig length. For each sample, they evaluate the best pairs of bins between a list of predicted bins *P* and a list of ground truth bins *T* . In Equations (12) and (13), *overlap*(*p, t*) is the sum of length of contigs in both the predicted bin *p* and the ground truth one *v*, while *size*(*b*) is the lengths sum of contigs in bin *b*. The F1 measure is the harmonic mean between *Prec* and *Recall*.

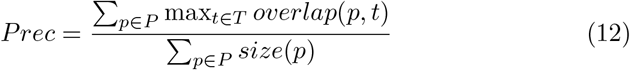

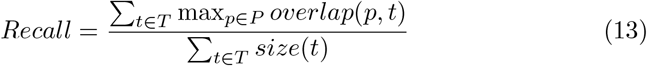

The second evaluation measures have been introduced in the plasmid binning evaluation tool `PlasEval` [11]. For each sample, `PlasEval` provides the costs associated to the minimum number of cut and join operations required to transform the predicted bins into the ground truth ones. The join and cut costs are linearly composed with the costs of extra predicted contigs and missing ground truth contigs to define a dissimilarity score, normalized between 0 and 1. In the next sections, we set the alpha `PlasEval` parameter to 0.5. While we aim maximizing the F1 score, we aim to minimize the dissimilarity score.

These measures are relevant to evaluate a binning result. Indeed, the binning results in a list of bins that may share some contigs. Consequently, classical clustering or partitioning evaluation measures do not handle this list. Furthermore, the adapted F1 and the dissimilarity measures are not affected by true chromosomal contigs that do not belong to any predicted bins.

### 3.3 Results

On the 839 samples, we keep those for which `PlasEval` returns an evaluation for each of the 6 methods (591 for the computation of the F1, 581 for the computation of the dissimilarity scores). Causes of the absence of an evaluation are explained in Appendix C.1.

For the best prediction-ground truth pairing evaluation, Figure 3 shows that `PlasBin-HMF` is slightly better (mean 0.61, median of 0.78, before `MOB-recon`, with 0.58 and 0.76), however the filtered version is the best (respectively 0.66 and 0.85, with a higher Q1 of 0.30 versus 0.07 for `MOB-recon`).

**Fig. 3.**
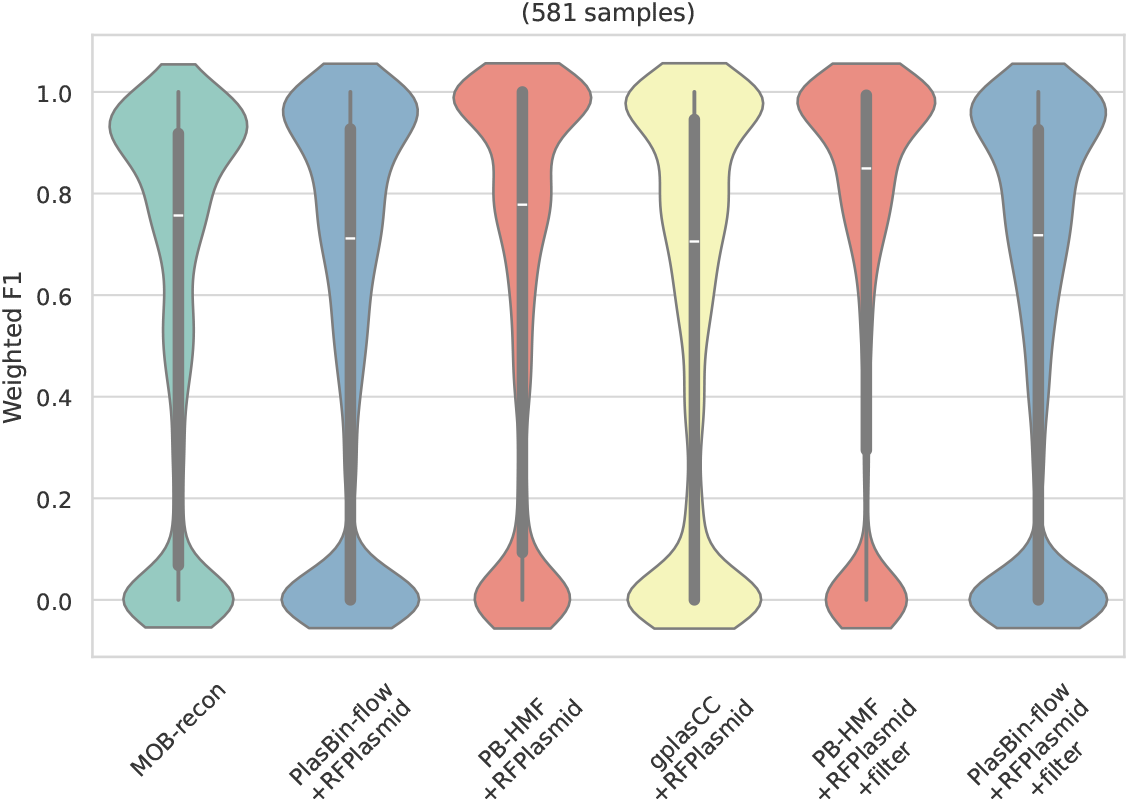
Weighted F1 scores

Figure 4 shows the dissimilarity scores distributions. `PlasBin-HMF` and its filtered version are the best (mean and median equal to 0.3 and 0.27, versus 0.39 and 0.3 for `gplasCC`), with Q1 quartile that reaches 0 (0.05 for `gplasCC`) and Q3 0.44 (0.66 for `gplasCC`). However, the conclusions vary depending on the sequenced bacteria species. `PlasBin-HMF` (especially filtered) keeps the best place for the *Staphylococcus aureus, Acinetobacter baumannii* and *Pseudomonas aeruginosa* species. While `MOB-recon` is not using an assembly graph, it is still efficient for *Escherichia coli* species for which it has been originally developed.

**Fig. 4.**
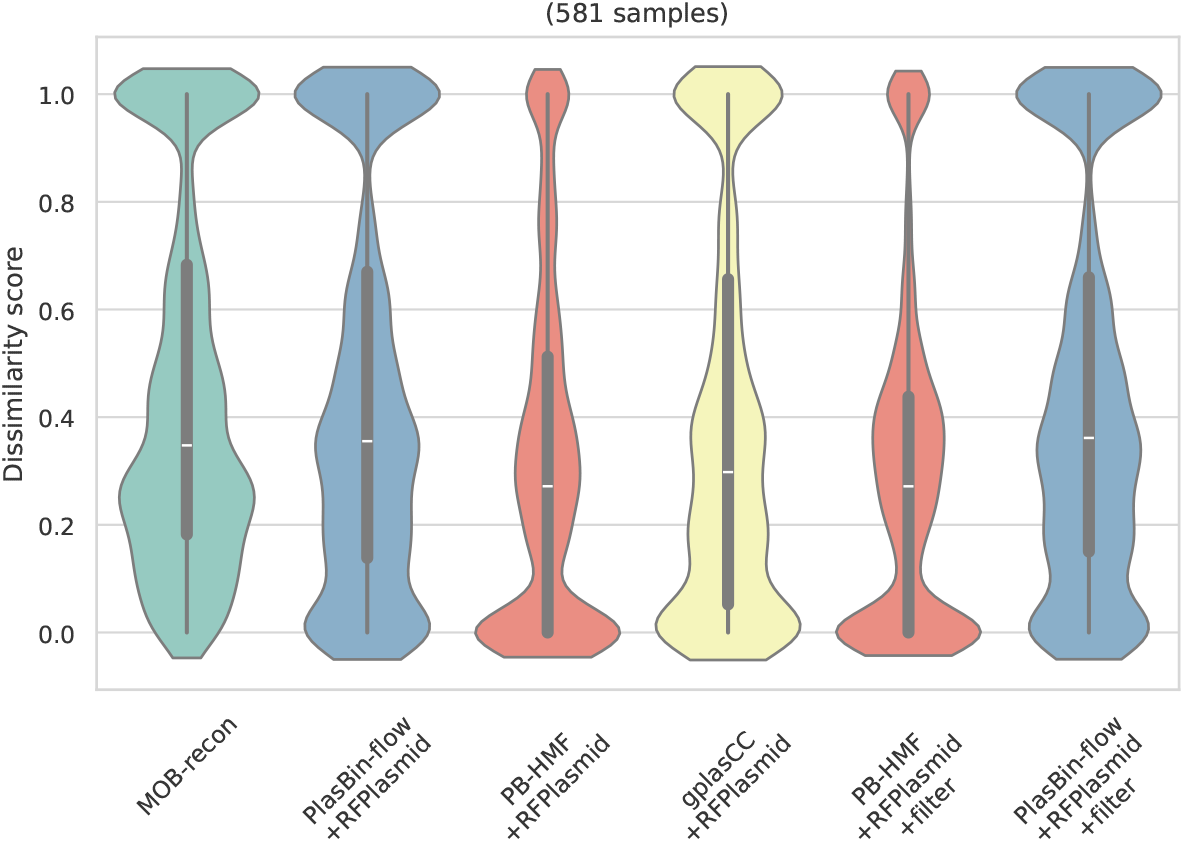
Dissimilarity scores

**Fig. 5.**
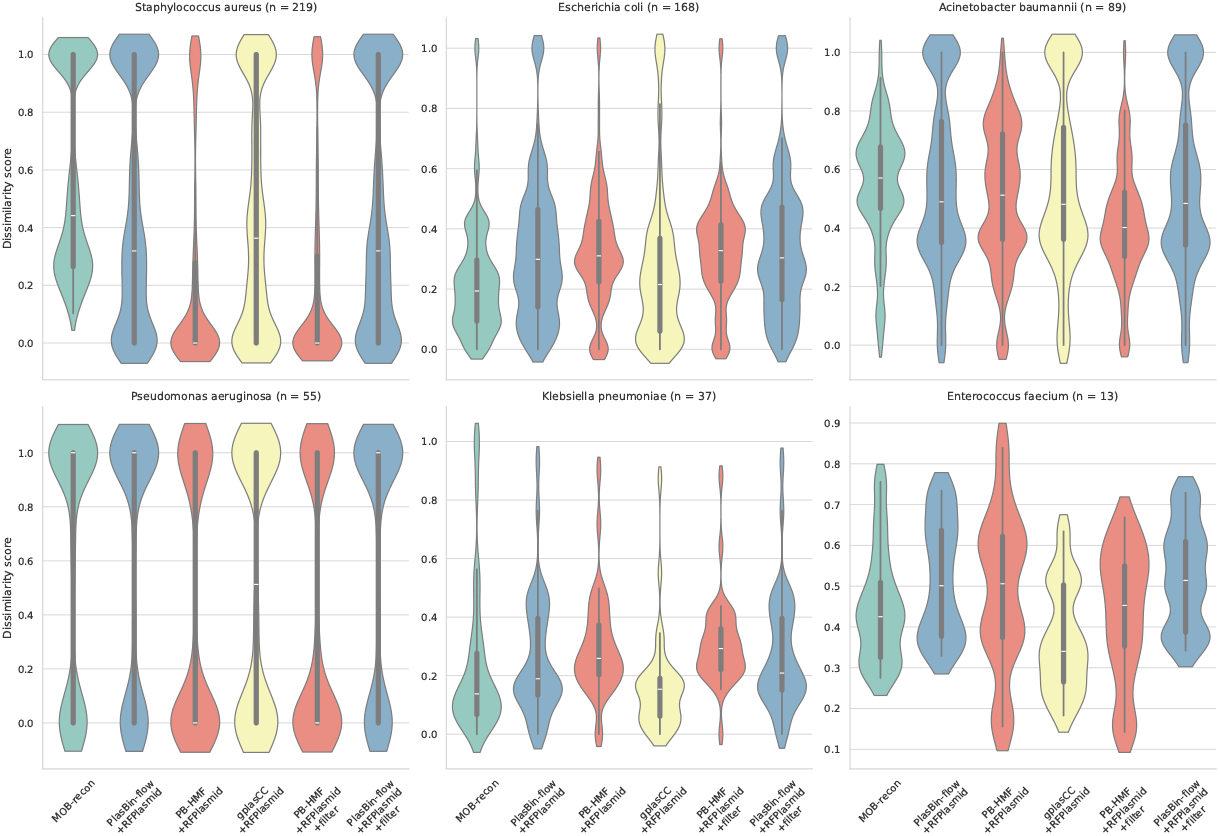
Dissimilarity scores per species.

One of the main hypothesis on the encouraging results of `PlasBin-HMF` is its high recall (and low missing contigs costs, Figures S5 and S7) combined with a moderate raise of the cut costs (the cost of splitting a bin that corresponds to several ground truth bins, Figure S8). On the one hand, we penalize the plasmidic coverages not consumed by the flows (Equation (10)). On the other hand, we search for circuits in the assembly graph, and due to the flow conservation constraints (Equation (3)), if two plasmids share contigs and their circularity is preserved, our model artificially tends to merge the two circular flows in one bigger circular flow. In that case, what was a shared contig between two plasmids is now a repeat in the artificial plasmid containing the two true ones. Although we have this bias by design, the plasmid subgraphs, while still connected, appear sufficiently dispersed in the assembly graphs such that we do not merge them. In other words, the assembler `Unicycler` correctly untangled the shared subsequences between the plasmids.

## 4 Conclusion

In this paper, we present the first hierarchical multi-flow approach to address the problem of recovering an unknown number of plasmid bins from a short-read assembly graph. Our methodology introduces a novel plasmid binning strategy that exploits the circular structure of most plasmids. We model the binning task as a combinatorial optimisation problem: we use a mixed-integer linear programming (MILP) formulation to search for a fixed, yet flexible, number of circular flows. Indeed, such number is increased during the process of computing flows. In a second step, we also consider partially circular flows to account for cases where missing links in the data disrupt plasmid circularity. To the best of our knowledge, `PlasBin-HMF` is the first method that simultaneously searches for multiple circular plasmids. We present experimental results comparing `PlasBin-HMF` with state-of-the-art binning tools, showing that our approach achieves the best performance on most bacterial samples.

Here we compared the exclusively binning tools (`gplasCC`, `PlasBin-flow` and `PlasBin-HMF`) with a classifier, `RFPlasmid` . Future comparisons would require us to use different classifiers, such as `plasmidCC`, the default classifier in `gplasCC` . Overall, the current results indicate that `PlasBin-HMF` has strong potential to substantially improve upon existing binning tools. However, several directions remain for further enhancing the model.

One of the main limitations of our approach is an inherent bias toward merging two circular plasmids when they share some contigs. One possible solution is to extend the MILP formulation with constraints on specific genetic content that should not be duplicated within a single plasmid. Such constraints must be carefully selected based on the plasmid genomics literature. Plasmid typing, which typically follows binning, could provide useful information for defining these constraints.

Alternatively, without introducing additional biological constraints and in order to keep the model minimalist, we could limit the amplitude of copy numbers (i.e., explained coverage variation) within each bin subgraph. This may help remove short, low-coverage chromosomal contigs that connect two plasmids and lead the model to incorrectly merge them. However, such a constraint may be less effective when separating plasmids with similar copy numbers. Similar challenges also arise in the haplotype assembly problem.

Another important direction concerns the number of flows in the multi-flow formulation, which corresponds to the unknown number of plasmids in the sample. Since this number is not known a priori, we currently implement a greedy iterative optimisation procedure to determine the appropriate number of flow functions. However, this strategy does not always scale well, as the branch-and-bound process may spend substantial time proving that the final multi-flow solution is not better than the previous one, leading to long runtimes before reaching an optimality certificate. In future work, rather than iteratively increasing the size of the multi-flow set, we aim to allow the MILP solver to directly decide the number of flows.

## Acknowledgments

This study was funded by the Simon Fraser University Project N001347 Grant and V. E., P.B and G.D.V. have received funding from the European Unions Horizon 2020 Innovative Training Networks programme under the Marie Skodowska-Curie grant agreement No. 956229. P.B and G.D.V have been funded by the Horizon Europe program under grant agreement No. 101160008 (FORGENOM II).

## Disclosure of Interests

The authors have no competing interests to declare that are relevant to the content of this article.

## A. Supplementary definitions

Figure S1 gives two ways of visualizing the assembly graph.

**Fig. S1.**
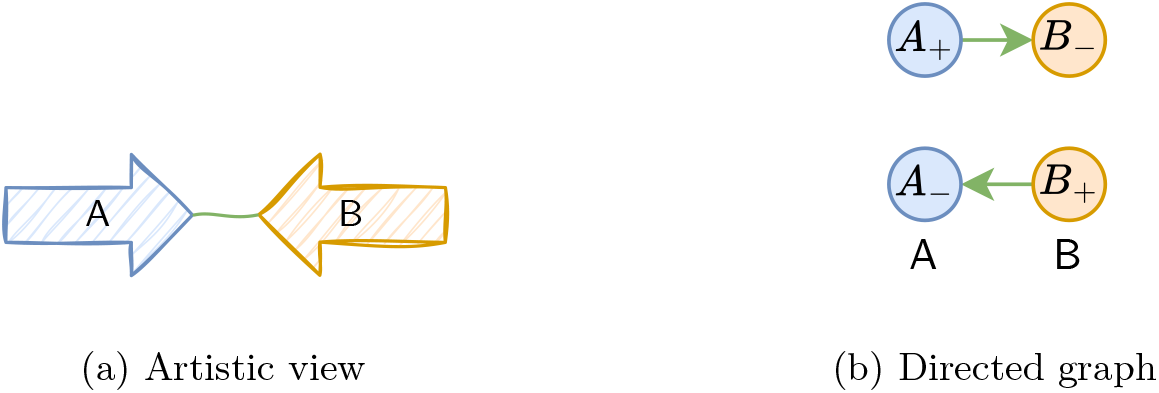
Assembly graph representations. The two subfigures represent the same information: two contigs A (in blue) and B (in orange), and a link between them (A+, B *−*) (in green). **(a)** The arrows are the oriented contigs, the green line is the link. **(b)** The directed graph structure is an explicit representation of the oriented contigs and their (oriented) links. Each vertex is one contig in a fixed orientation. One link and its reverse are the arcs.

## B Supplementary method

### B.1 Method overview

The multi-flow binning approach searches several bins with the same properties at the same time. The properties are of two types:

#### Topology

The bin is circular 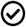, otherwise partially circular 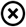

#### Seed constraint

Each bin must contain at least one seed 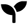, otherwise, it can be free of seeds 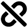

##### Algorithm B.1 Hierarchical Multi-Flow binning (HMF)

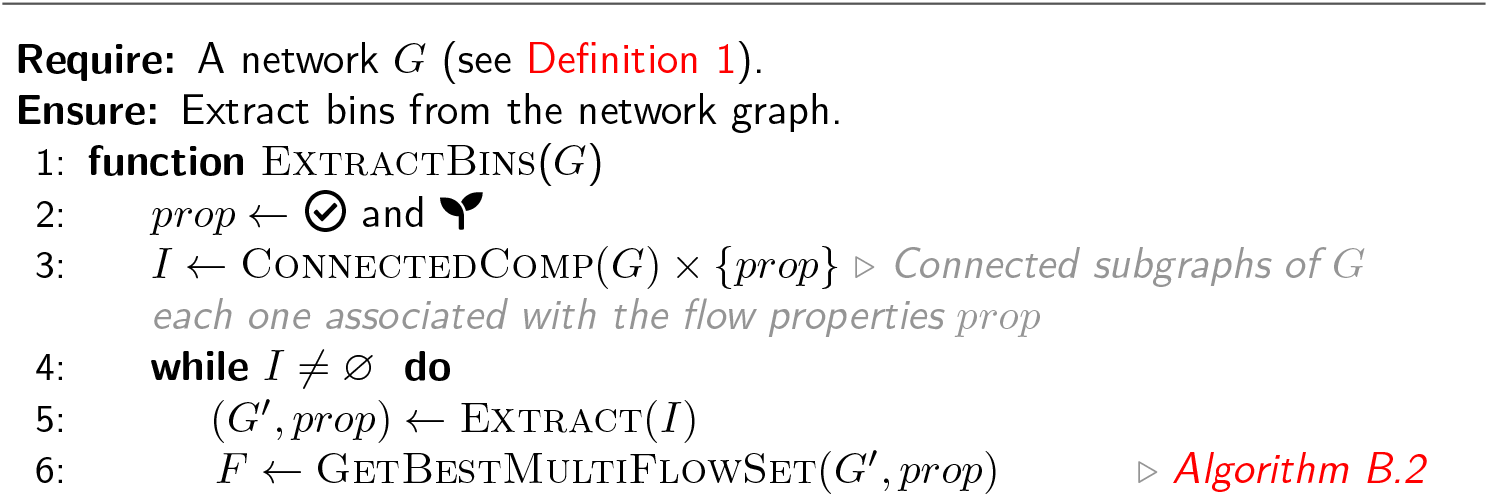
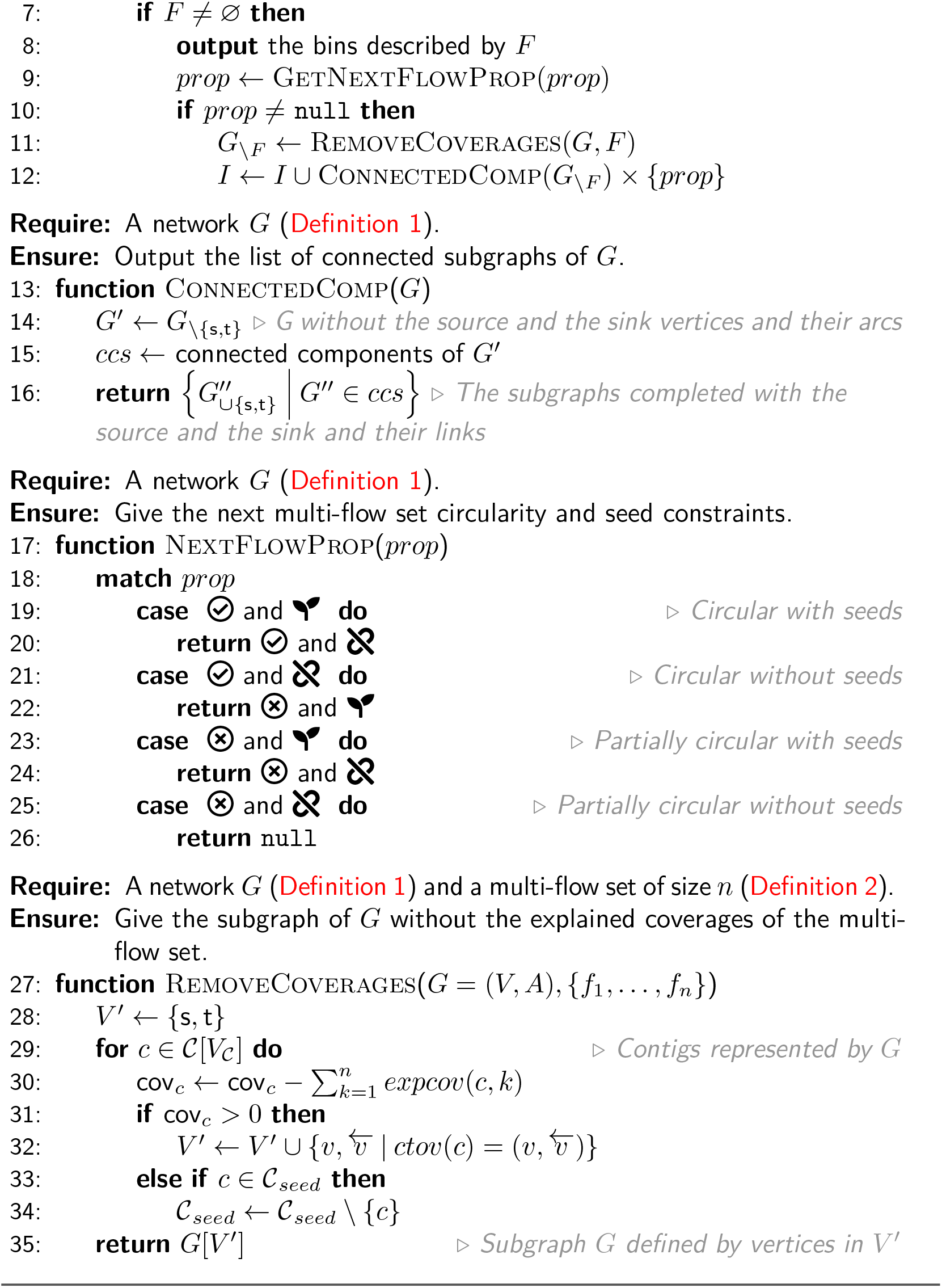

##### Algorithm B.2 Find the best multi-flow set with the best size

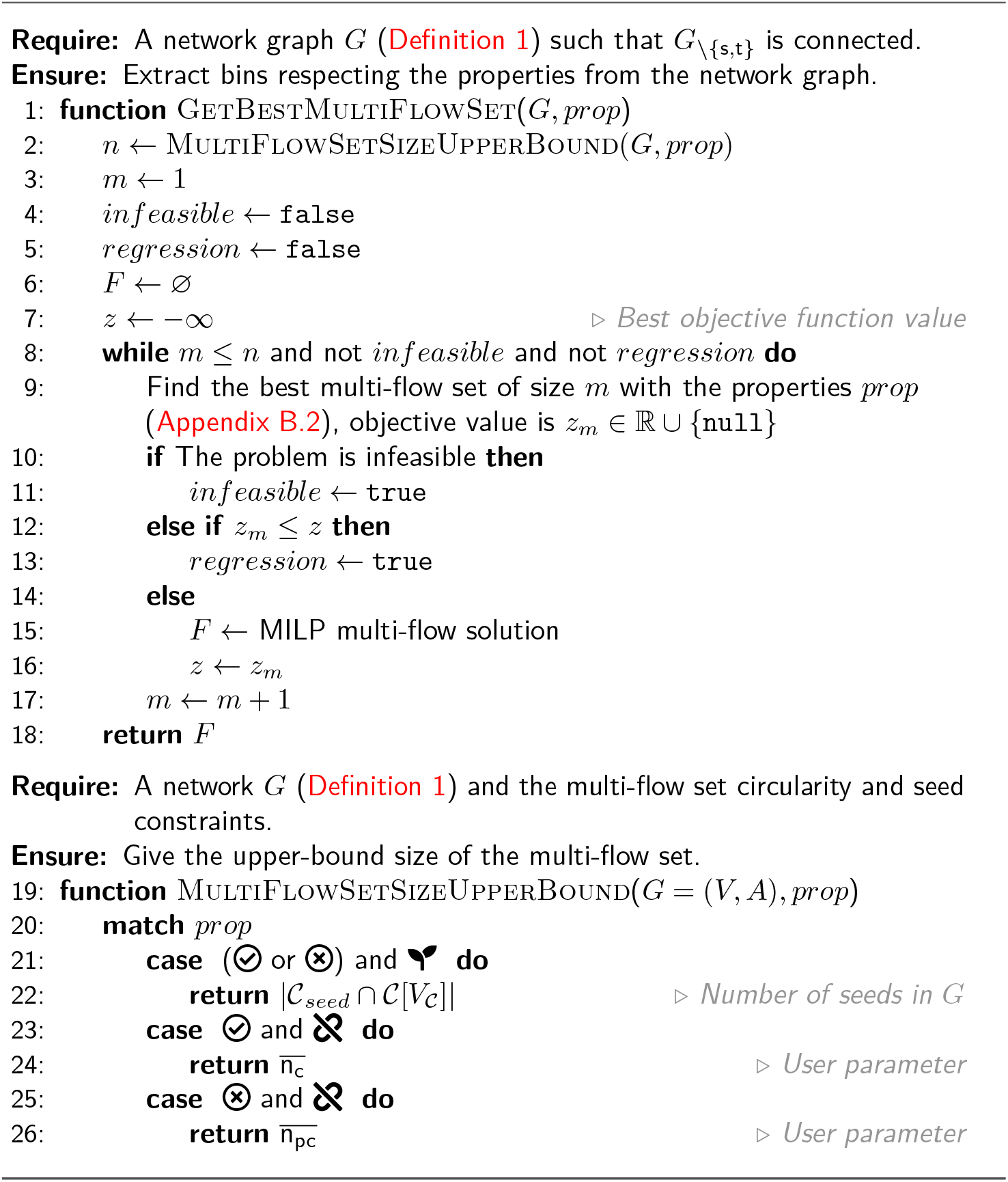

### B.2 Mixed Integer Linear Programming model

In this section we present the MILP model for the multi-binning flow approach, where one flow induces one bin. The MILP formulation consists in a set of constraints where the right inequality constant arm can change to make correspond the model to the flow/plasmid properties. Let *n* ∈ ℕ be the maximum number of bins we can or want to model, and *k* ∈ {1, …, *n*} thus representing the *k*^th^ flow. The changes of the constraints’ constant parts must cover the following states:

– Flow *k* is inactive 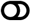 (no bin induced).
– Flow *k* is active 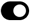 (a bin is induced), with the following properties:
  - it is circular 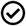 or partially circular 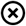;
  - it contains at least one seed 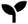 or can be free of seeds 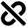

We associate a symbol to each state to easily identify to which constant constraint right arm it corresponds to. The flow property symbols 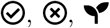 and 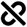 imply the flow to be active 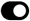 so we do not repeat the last symbol. The additional symbol 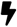 means the constraint only applies for performance optimization.

In the following sections, by *A*_*ℒ*_ we denote the set of link-arcs such that each link-arc corresponds to a link in ℒ.

**Variables** Table S1 lists all the variables used in the MILP model.

**Table S1.**
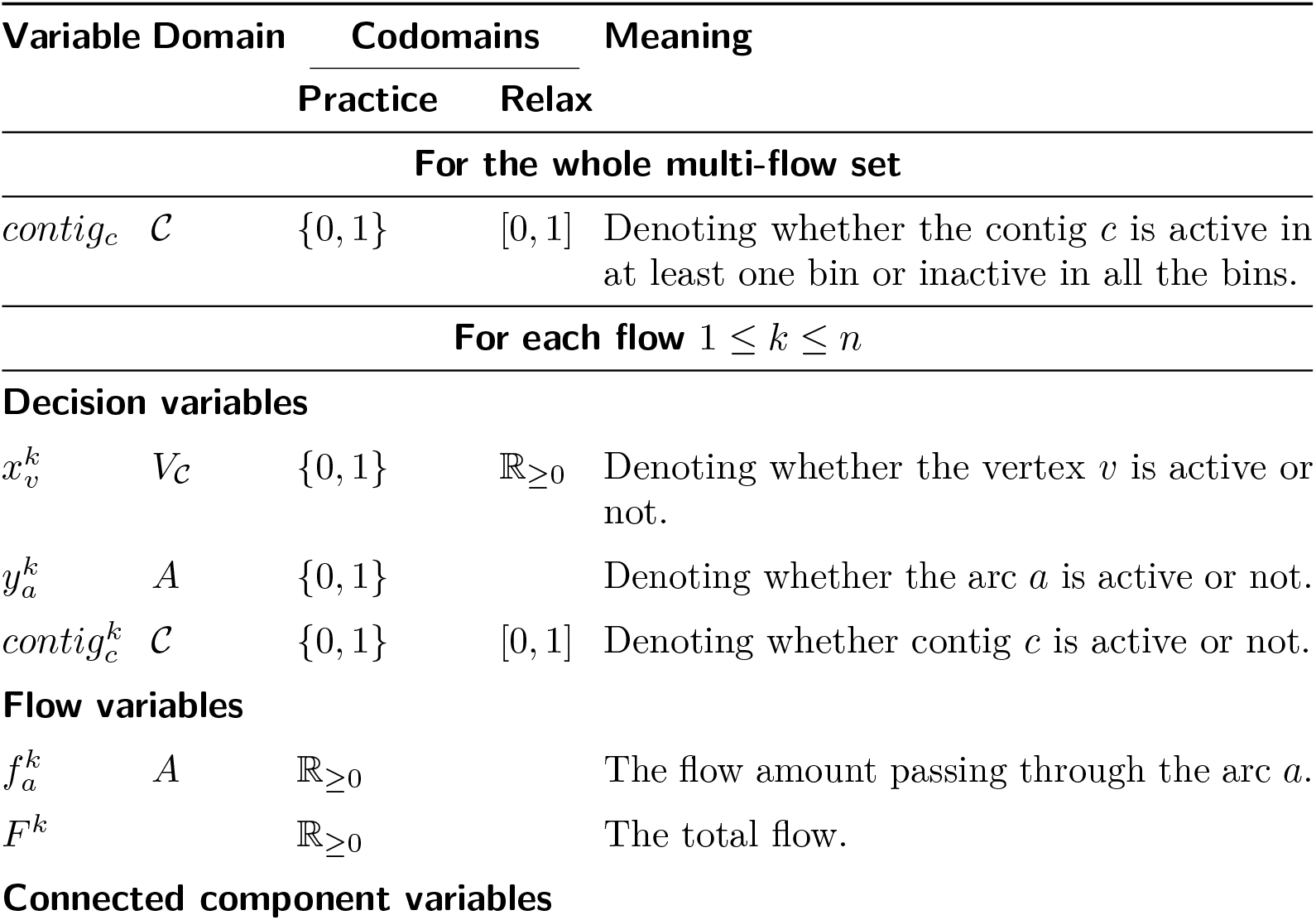
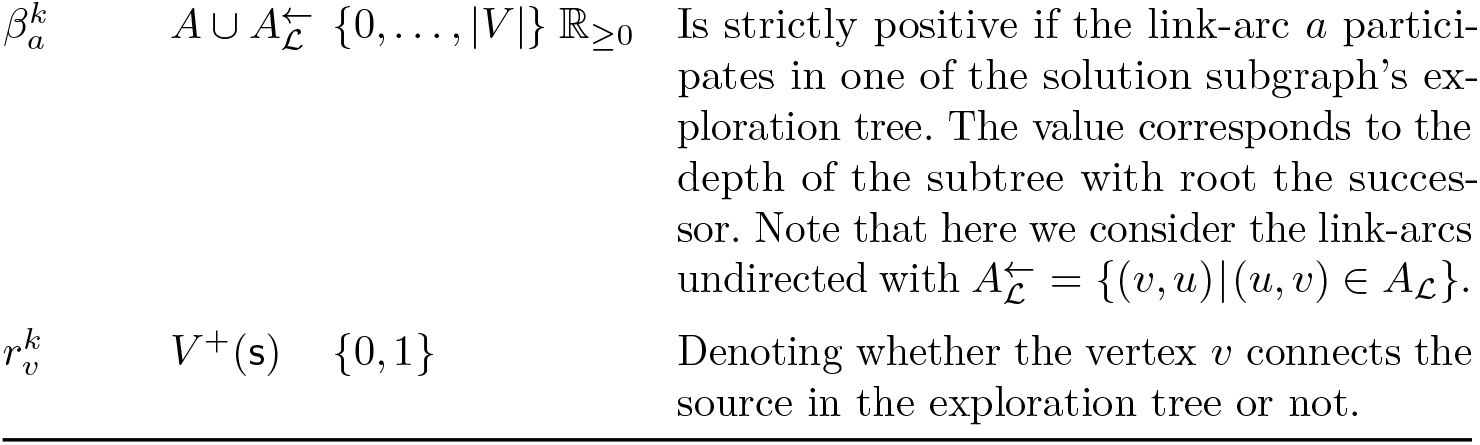
MILP variables. Each of the following variables participates in at least one of the MILP model. Some of them participate in all the model, others are necessary for only one model. For the ease of read, we categorize the variables in three categories. Each section describing a model precise which variables are participating for each category.

#### Constraints for the whole bins

This section gives the constraints for all the bins without distinction. The next section provides the constraints for one bin.

The cumulative incoming flow among all the bins must not exceed the coverage of the contig:

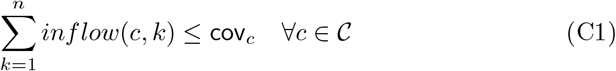

Where

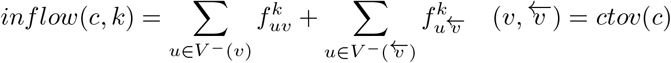

Here we define the behaviour of the variable *contig*_*c*_ equals to 1 if the contig is active in at least one bin, otherwise 0:

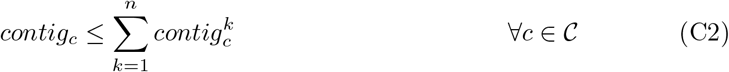

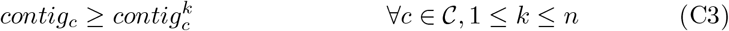

When the number of flows to be active is greater than one (2 ≤ *m ≤ n*, the explanation score (*ExplanationScore*_*m*_) penalized by the flow cost must be greater or equal to the previous best objective value ExplanationScore^*^ (see Appendix B.2).

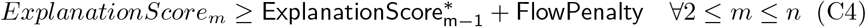

Where FlowPenalty = <u>L</u> · <u>cov</u> corresponds to the penalty cost of explaining the data with an additional flow. Remind that <u>L</u> is the minimum cumulative contig length, and <u>cov</u> is the minimum amount of flow passing through a link-arc. The term is multiplied by 1 which corresponds to the best plasmidness.

When the number of bins to be active is greater than one, we define a plasmidness score order between the bins to avoid bins permutation:

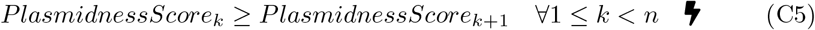

#### Constraints for one specific flow

Let *k* ∈ *{*1, …, *n}* be the flow number.

*Decision variables relationships* A contig is active if, and only if at least one of its corresponding vertices is active:

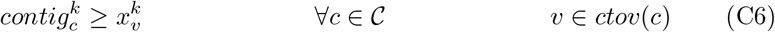

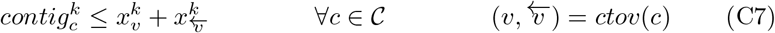

The vertices involved in an active arc must also be active:

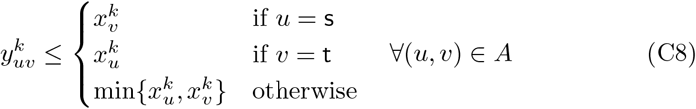

An active vertex implies at least one active arc incoming to it:

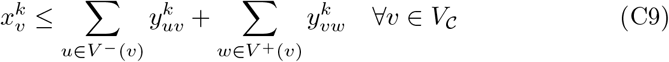

*Flow constraints* for each contig vertex, the incoming flow equals the outgoing flow:

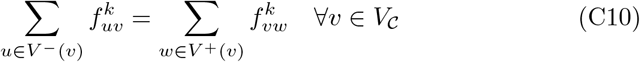

The total flow equals both the source outgoing flow and the sink incoming flow:

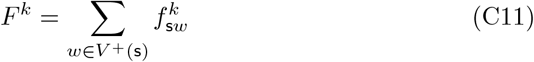

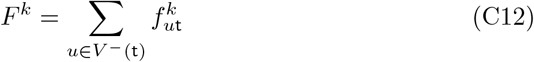

Each arc flow cannot exceed the coverage of the contigs involved. Also, the flow of an active arc is lower bounded by <u>cov</u> > 0, except the source and sink active arcs in the circularity context, where their flow equal 0:

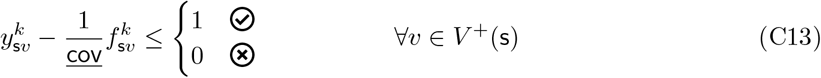

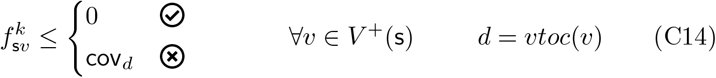

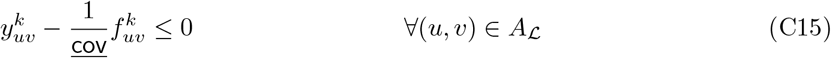

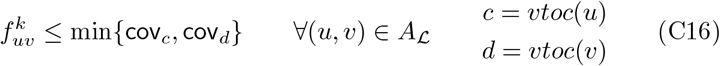

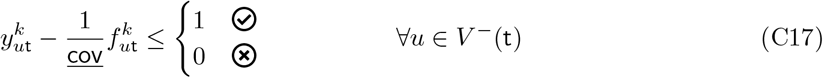

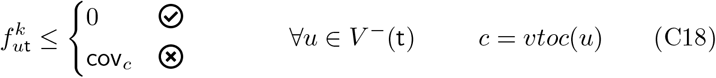

The cumulative incoming flow cannot exceed the coverage of the contig:

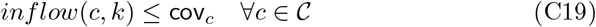

*Connectivity constraints* To avoid the solution graph without the source and the sink (the intermediate solution graph) to have several connected components, we inspire the Connected Intermediate Flow problem in [7] and adapt the constraints to the nature of our network. Modelling the connectivity implies describing an exploration tree.

An active arc can participate in the tree. Also, an active link-arc in *A*_L_ can reversely participate in the tree 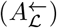:

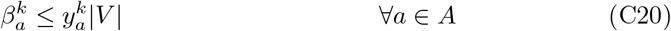

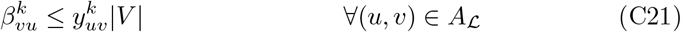

Note that in Constraint C21, 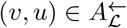

In the circular case, we do not need to model the reverse link-arc in the tree:

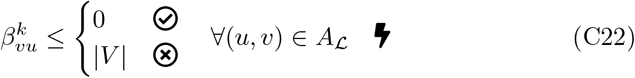

To define the whole tree, we must define each subtree each arc (or reversed link-arc) defines. We must be careful of the validity of the constraints for the source and the sink when the bin is inactive.

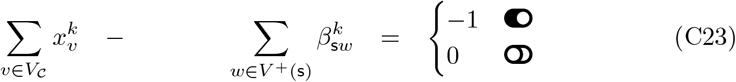

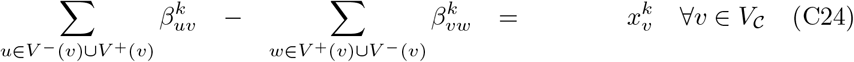

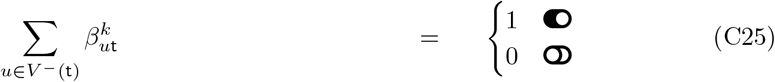

Only one contig vertex connects the source in the tree. It is a key constraint to model the connectivity of the intermediate solution graph:

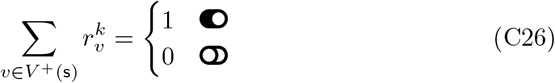

When the bin is circular and must contain a seed, as the benefit of the performance, we only authorize the seed vertices to connect the source in the graph, so in the tree:

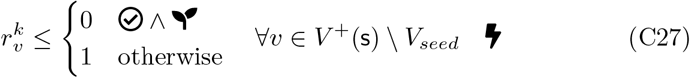

Only the contig vertex chosen to be the root can connect the source in the tree:

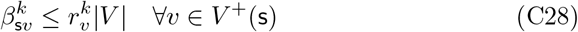

Several contig vertices can connect the source in the solution graph. But only one (the root) connects the source in the tree. For performance purposes, we can arbitrarily give an order on the root choice.

Let **v**_s_ to be an arbitrary vector containing once all the vertices connected to the source (order of *V* ^+^(s)). Then:

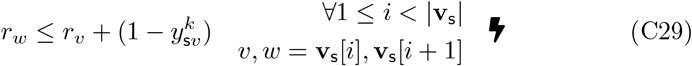

Thus, the first vertex in **v**_s_ that connects the source in the solution graph is the one which connects the source in the tree.

*Plasmid property* The cumulative contig length is above a given threshold:

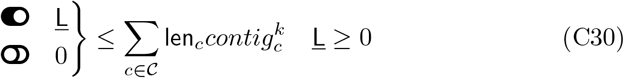

Typically, we choose <u>L</u> = 1000.

The incoming flow weighted plasmidness is above its *γ* coefficient dynamic mean:

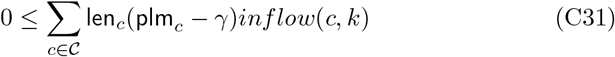

Typically, we choose *γ* = 0 to ensure the plasmidness of the bin to be positive.

*To be circular or to be partially circular* The circular bin is defined by only one source-arc, respectively one sink-arc, while the partially circular bin can be defined by several source-arcs, respectively several sink-arcs:

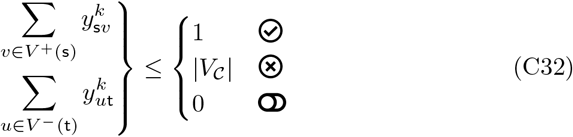

When the bin is circular, each active contig must follow and precede another contig:

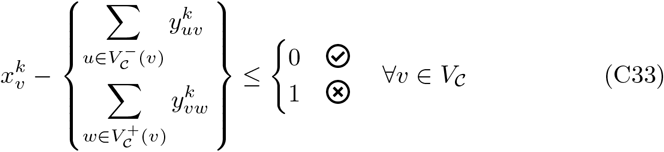

When the bin is partially circular, an active contig that connects the source must not follow another contig:

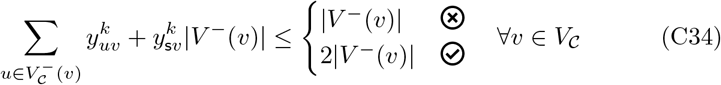

Analogously to the previous constraint, when the bin is partially circular, an active contig that connects the sink must not precede another contig:

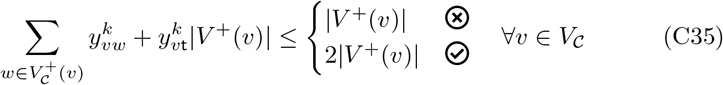

When the bin is circular, we can fix the same vertex to connect both the source and the sink:

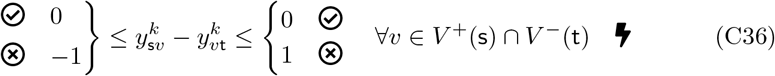

*Model whether the bin must have a seed or can be free of seeds* Set the minimum number of seed contigs in the bin:

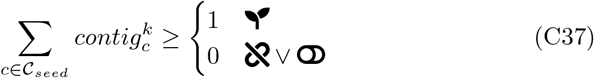

When the bin is circular, and must contain a seed, we can forbid the non-seed vertices to connect the source and the sink:

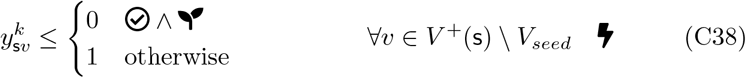

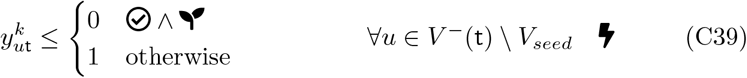

**Objectives** The MILP objective follows Definition 6.

**Definition 8 (MILP objective)**..

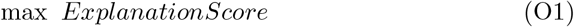

*Where*

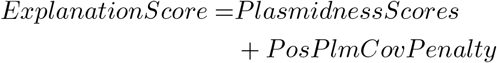

*such that:*

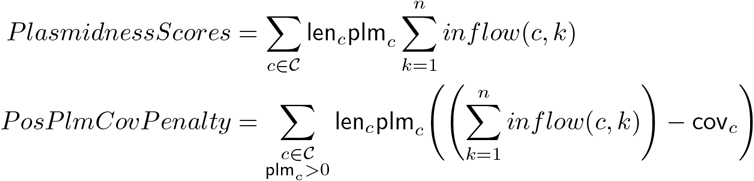

## C Supplementary results

We select the same 1241 bacteria sequencing projects samples from various bacteria species on the NCBI as in [12]^3^. Among the 1241 samples, 836 have a ground truth without unlabelled hybrid contigs. Of the 836 samples, a method can fail to return bins, either because it ends without error and with no bins, or because it reaches the time limit before having the time to output any bin. Even if a tool outputs bins, `PlasEval` dissimilarity computation (`comp` command) can fail to return an evaluation because the branch and bound it relies on has reached an upper-bound number of iterations, or because the script reaches the `sbatch` time limit (3 hours). We selected the samples for which every tools we compare output bins and have a `PlasEval` evaluation. In the following, for each figure, the numbers under parenthesis are the numbers of samples for which there is a `PlasEval` evaluation.

### C.1 Evaluation presence count

Note that in this section, the method order is not respecting the one given in the violin plots.

**Fig. S2.**
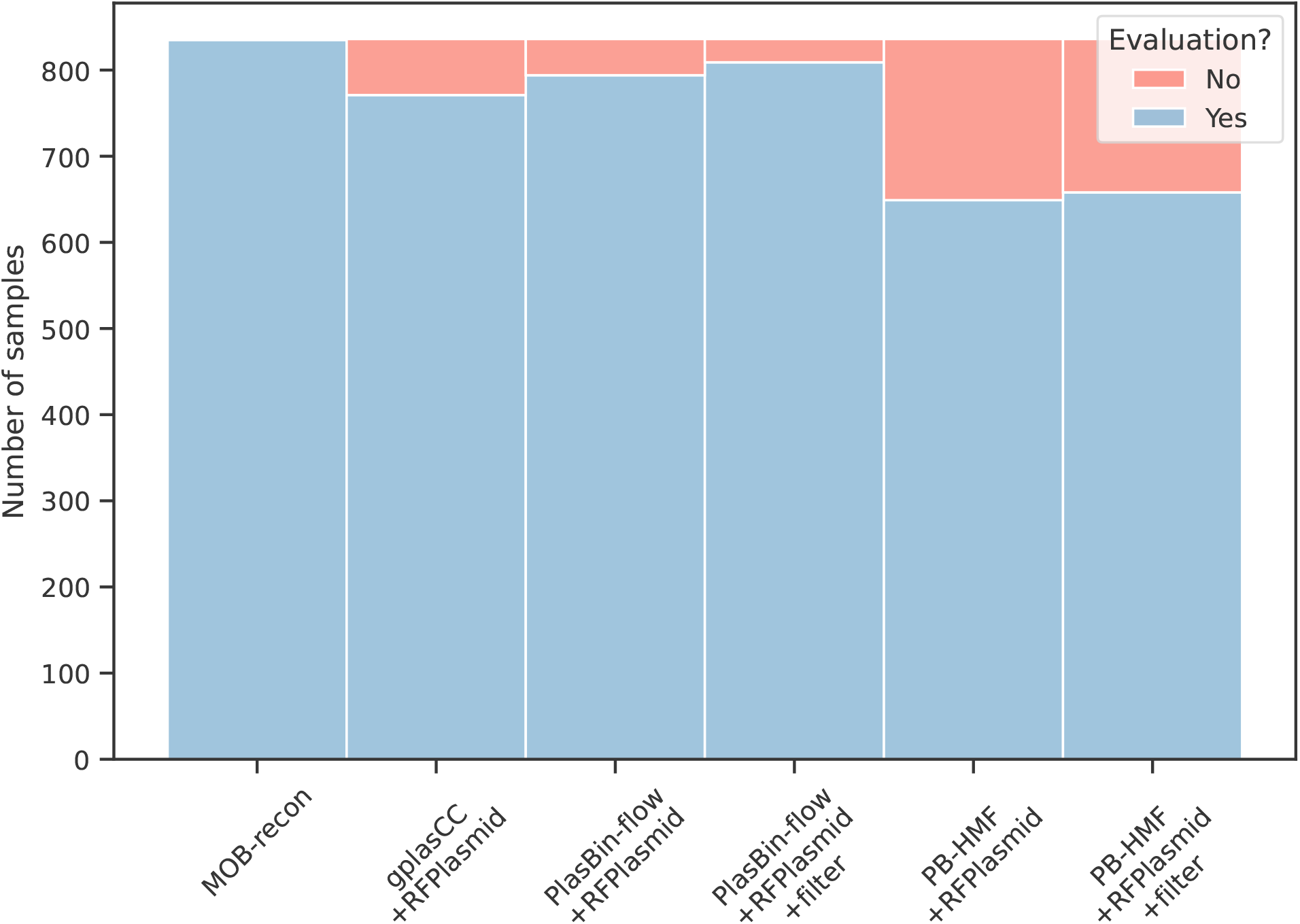
Number of `PlasEval` dissimilarity evaluations.

**Fig. S3.**
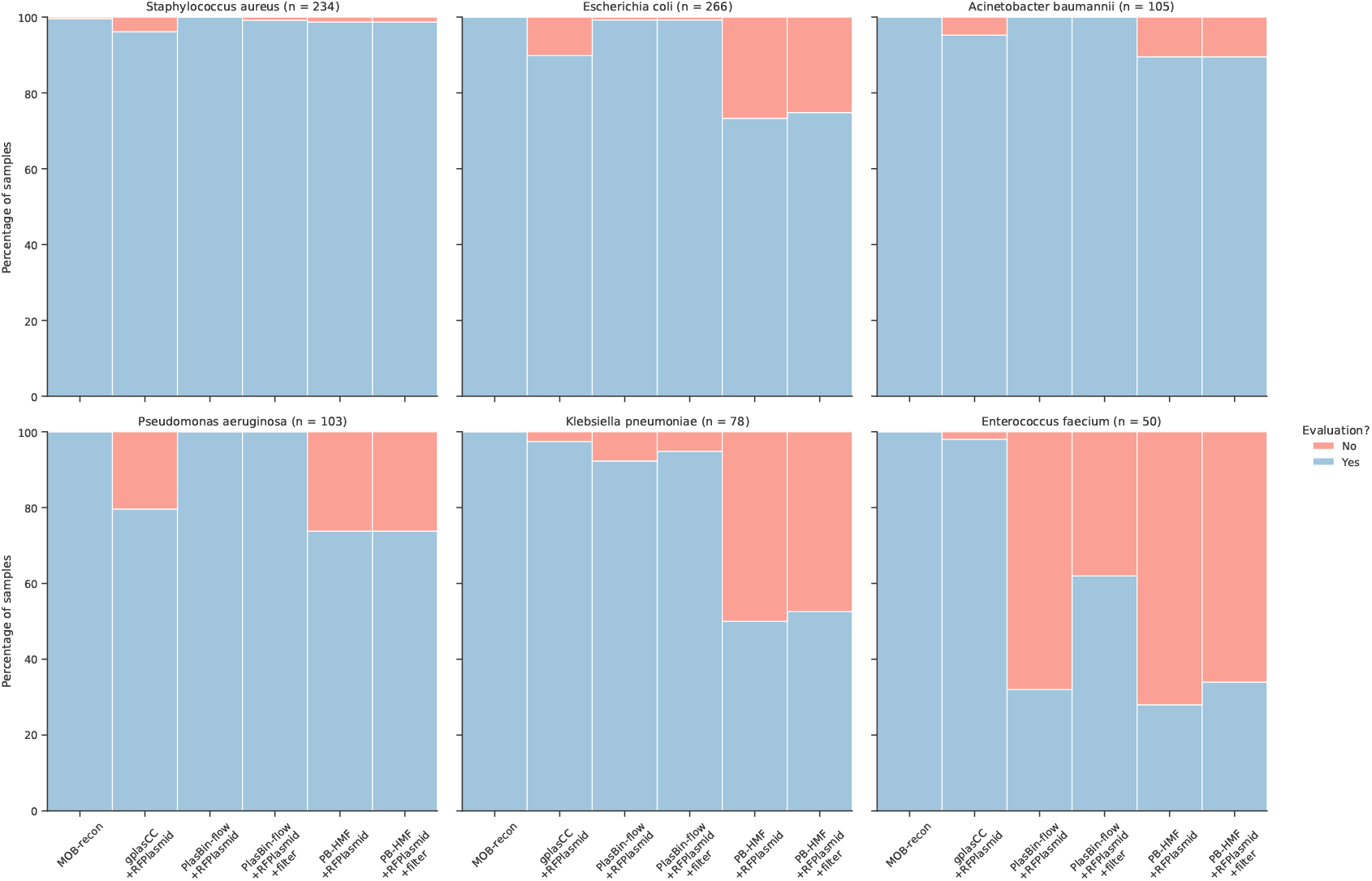
Number of `PlasEval` dissimilarity evaluations per species.

### C.2 Precision, recall and F1

Precision (Figure S4), recall (Figure S5) and F1 score (Figures 3 and S6 and table S2) are obtained with `PlasEval eval` command.

**Fig. S4.**
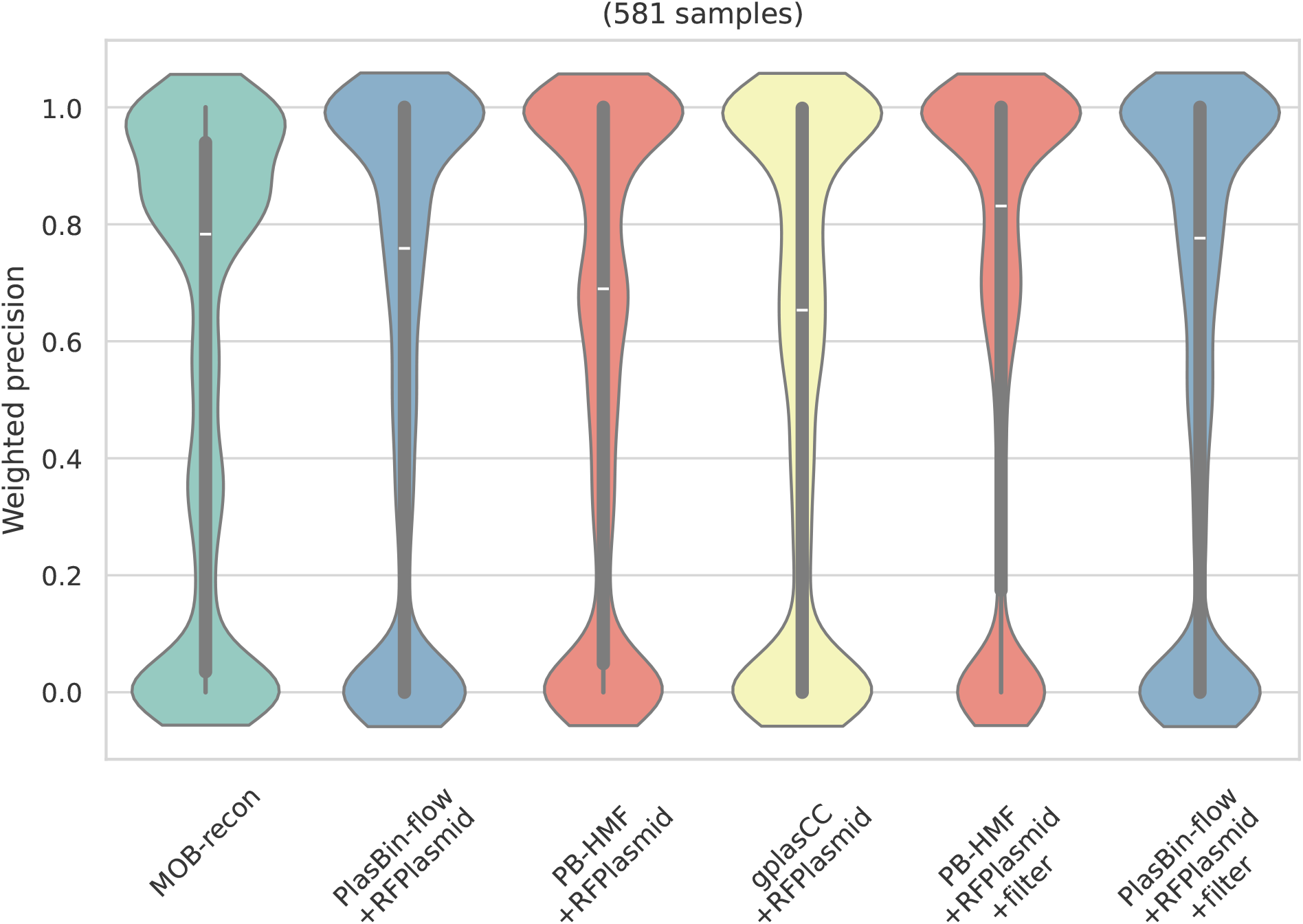
Precision.

**Fig. S5.**
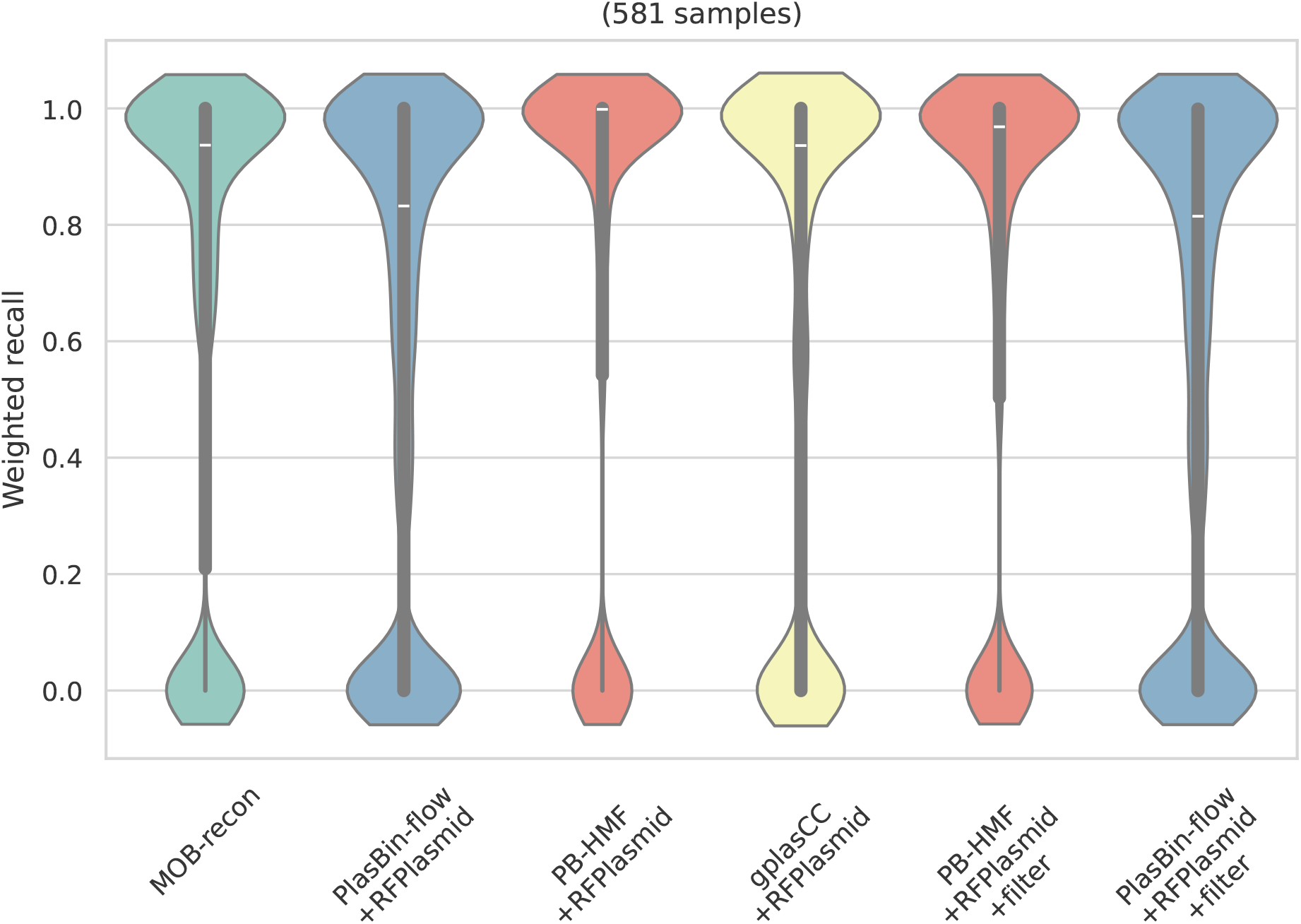
Recall.

**Fig. S6.**
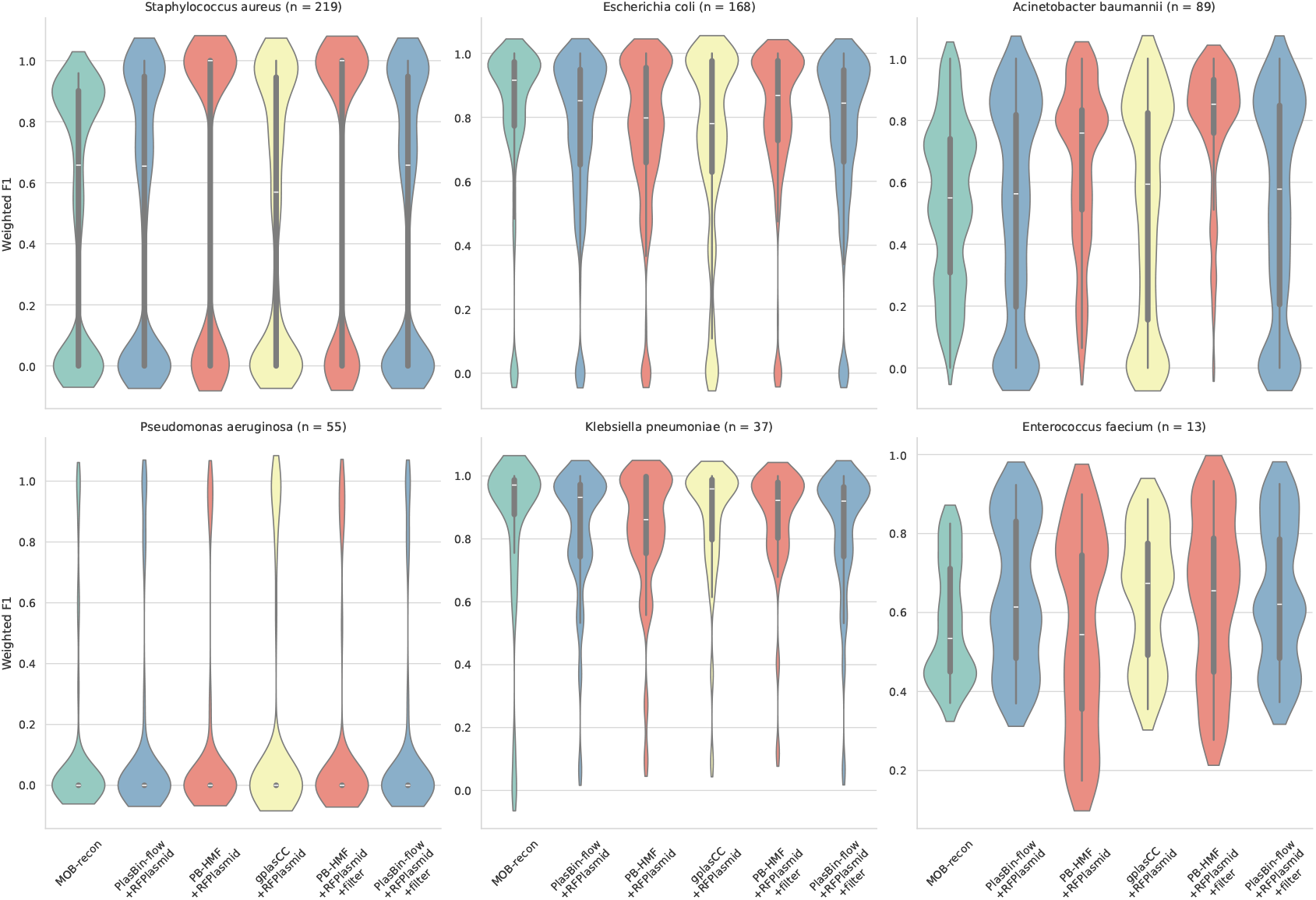
F1 per species.

**Table S2.**
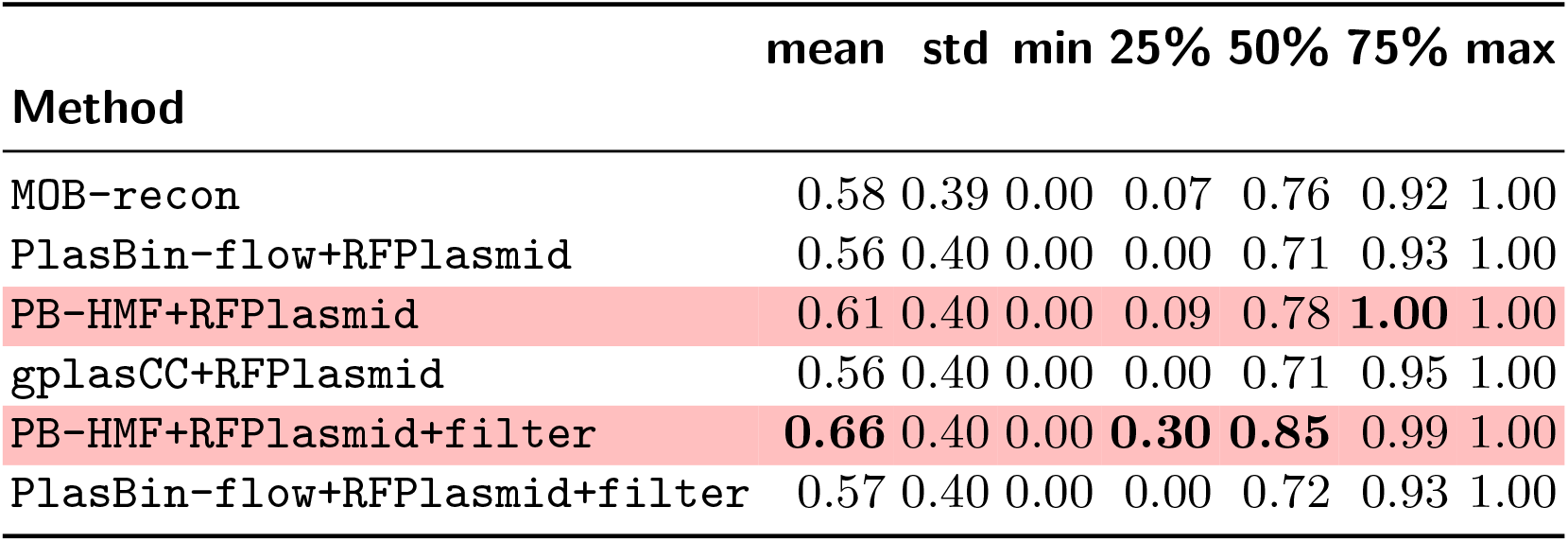
F1 statistics.

| Method | mean | std | min | 25% | 50% | 75% | max |
| --- | --- | --- | --- | --- | --- | --- | --- |
| MOB-recon | 0.58 | 0.39 | 0.00 | 0.07 | 0.76 | 0.92 | 1.00 |
| PlasBin-flow+RFPlasmid | 0.56 | 0.40 | 0.00 | 0.00 | 0.71 | 0.93 | 1.00 |
| PB-HMF+RFPlasmid | 0.61 | 0.40 | 0.00 | 0.09 | 0.78 | **1.00** | 1.00 |
| gplasCC+RFPlasmid | 0.56 | 0.40 | 0.00 | 0.00 | 0.71 | 0.95 | 1.00 |
| PB-HMF+RFPlasmid+filter | **0.66** | 0.40 | 0.00 | **0.30** | **0.85** | 0.99 | 1.00 |
| PlasBin-flow+RFPlasmid+filter | 0.57 | 0.40 | 0.00 | 0.00 | 0.72 | 0.93 | 1.00 |

### C.3 Dissimilarity scores

The dissimilarity score (Figures 4 and 5 and table S3), a linear composition of cut (Figure S8), join and extra and missing contigs costs (Figure S7), is obtained with `PlasEval comp` command.

**Fig. S7.**
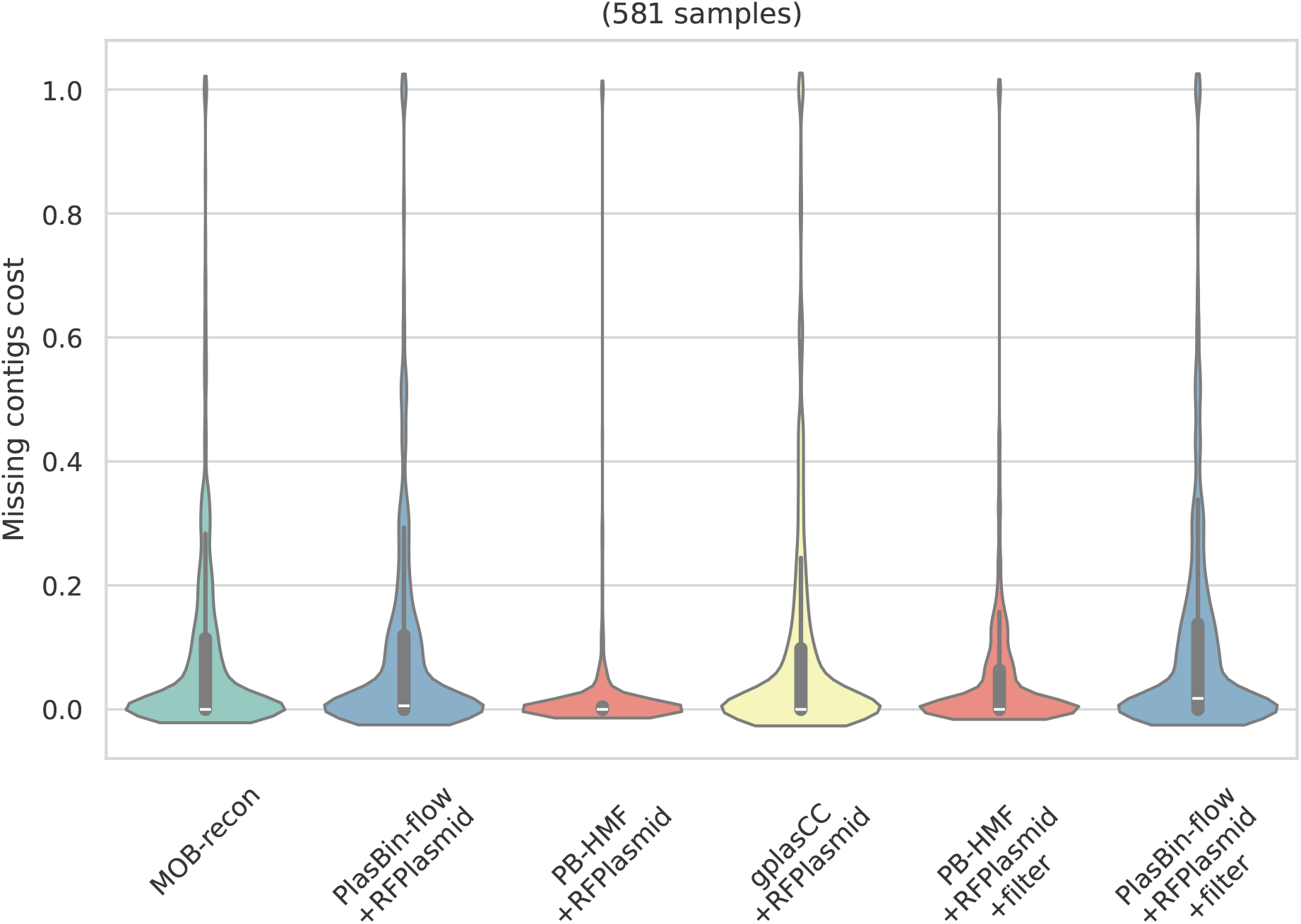
Missing contigs costs.

**Fig. S8.**
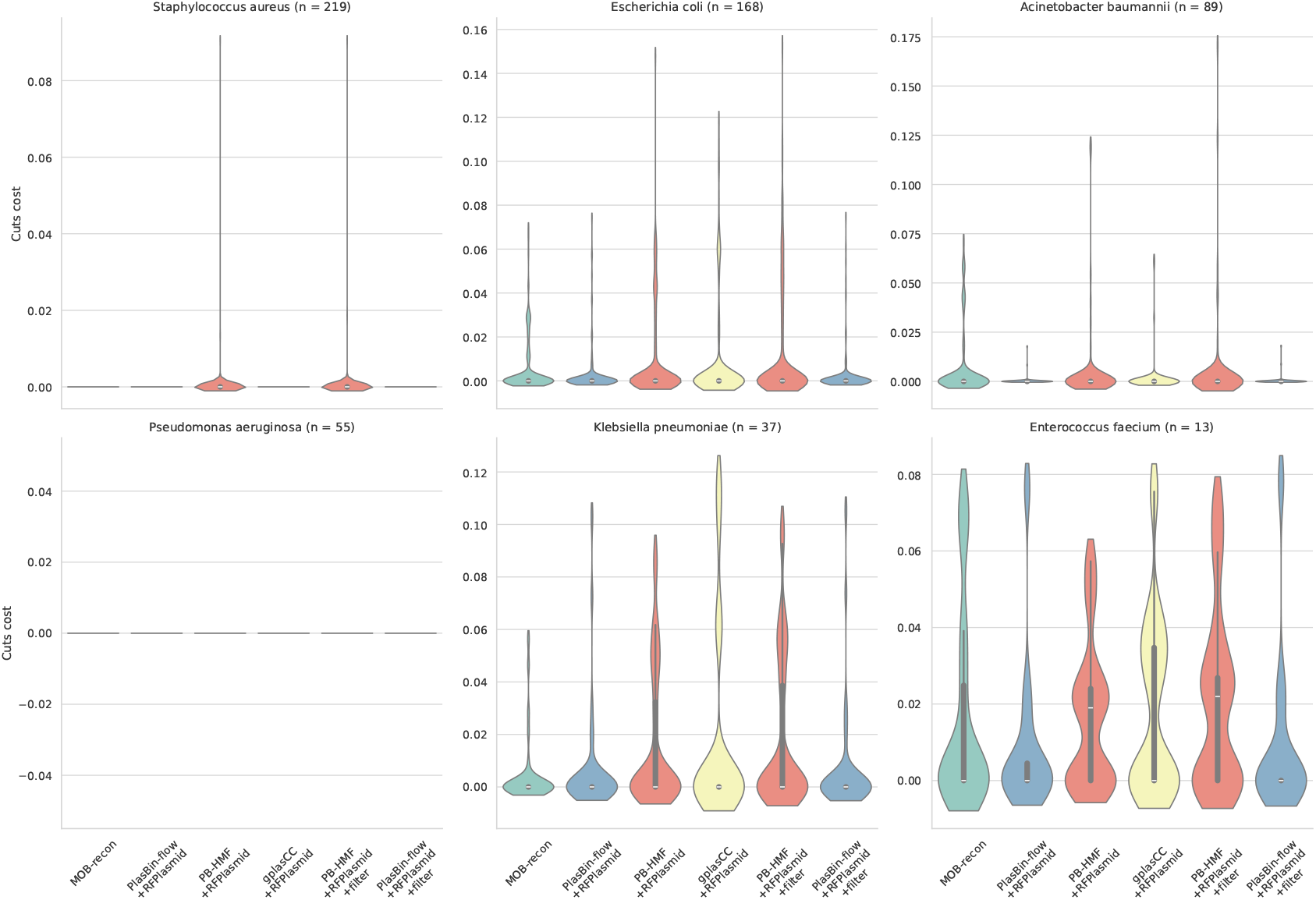
Cut costs per species.

**Table S3.** Dissimilarity statistics.

| Method | mean | std | min | 25% | 50% | 75% | max |
| --- | --- | --- | --- | --- | --- | --- | --- |
| MOB-recon | 0.45 | 0.34 | 0.00 | 0.18 | 0.35 | 0.68 | 1.00 |
| PlasBin-flow+RFPlasmid | 0.44 | 0.36 | 0.00 | 0.14 | 0.36 | 0.67 | 1.00 |
| PB-HMF+RFPlasmid | 0.33 | 0.33 | 0.00 | **0.00** | **0.27** | 0.51 | 1.00 |
| gplasCC+RFPlasmid | 0.39 | 0.37 | 0.00 | 0.05 | 0.30 | 0.66 | 1.00 |
| PB-HMF+RFPlasmid+filter | **0.30** | 0.31 | 0.00 | **0.00** | **0.27** | **0.44** | 1.00 |
| PlasBin-flow+RFPlasmid+filter | 0.44 | 0.35 | 0.00 | 0.15 | 0.36 | 0.66 | 1.00 |

## Footnotes

1 *V*_𝒞_ = 𝒞 *×{*+, −*}* is an abuse of notation. Rigorously, one should define *V*_*𝒞*_ as a regular vertex set in bijection with 𝒞 *× {*+, −*}*. The same remark holds for the arcs where one should define a set of link-arcs *A*_ℒ_ in bijection with ℒ.

2 Pending the submodule becoming a stand-alone programme, `PlasBin-HMF` can be launched via the command `pangebin asm-pbf hmf --help` to output a file in the same format as `PlasBin-flow`.

3 We reached out to the authors of the recent plasmid classification and binning tools benchmarking article [22] to gain access to their data, but have not received any response as of now.

## Notes

https://github.com/AlgoLab/pangebin

### Competing Interest Statement

The authors have declared no competing interest.

